# Ursolic acid improves the bacterial community mapping of the intestinal tract in liver fibrosis mice

**DOI:** 10.1101/2020.02.10.941955

**Authors:** Sizhe Wan, Chenkai Huang

## Abstract

Liver fibrosis often appears in chronic liver disease, with extracellular matrix (ECM) deposition as the main feature. Due to the presence of the liver-gut axis, the destruction of intestinal homeostasis is often accompanied by the development of liver fibrosis. The inconsistent ecological environment of different intestinal sites may lead to differences in the microbiota. The traditional Chinese medicine Ursolic acid (UA) has been proven to protect the liver from fibrosis. We investigated the changes in the microbiota of different parts of the intestine during liver fibrosis and the effect of UA on these changes based on high-throughput sequencing technology. Sequencing results suggest that the diversity and abundance of intestinal microbiota decline and the composition of the microbiota is disordered, the potentially beneficial Firmicutes bacteria are reduced, and the pathways for functional prediction are changed in the ilea and anal faeces of liver fibrosis mice compared with normal mice. However, in UA-treated liver fibrosis mice, these disorders improved. It is worth noting that the bacterial changes in the ilea and anal faeces are not consistent. In conclusion, in liver fibrosis, the microbiota of different parts of the intestines have different degrees of disorder, and UA can improve this disorder. This may be a potential mechanism for UA to achieve anti-fibrosis. This study provides theoretical guidance for the UA targeting of intestinal microbiota for the treatment of liver fibrosis.

## INTRODUCITON

Hepatic fibrosis often appears in chronic liver disease, with extracellular matrix (ECM) deposition as its main feature^1^. Proliferating connective tissue blocks the regeneration space of normal liver cells, causing damage to liver structure and function^2^. The continued development of liver fibrosis can lead to liver cirrhosis and even liver cancer, causing more than 500,000 deaths each year^3^. As an early pathological change, liver fibrosis has reversible characteristics and is therefore valued. However, the underlying mechanisms of liver fibrosis development have not been fully explored. It is currently believed that the activation of stellate cells is stimulated, and proliferation and migration are considered central events of liver fibrosis^4^. Activated HSCs lead to a reprogramming of liver metabolism, increased autophagy and increased parenchymal cell damage, leading to a loss of hepatic stellate cell (HSCs) retinoids, increased contractility, and amplification of growth factors and inflammatory signalling factors in the liver microenvironment, which in turn produces a large amount of ECM and promotes fibrosis occurrence and development, etc^5 6^.

The human gut is home to a microbial community of at least 1,000 species of bacteria, up to 10^14^ species of microorganisms^7^. The interaction of the host with the microorganism is critical to the normal physiological functions of the host, from metabolic activity to immune homeostasis^8-10^. Changes in gut microbiota are thought to be closely related to many diseases^11-13^.

There is a special anatomical and positional relationship between the liver and the intestine; that is, 75% of the blood in the portal vein is supplied by the intestine. This special relationship makes the intestines and the liver closely connected, and the two interact with each other, which is the “liver-gut axis”^14^. The liver regulates intestinal homeostasis through immune regulation, energy metabolism, and bile acid excretion^15 16^. Currently, disorders of the intestinal microbiota have been found in many chronic liver diseases^17-19^. As an important component of the Gram-negative bacterial wall, LPS can enter the blood through the damaged intestinal barrier and enter the liver through the gut-liver axis, inducing the liver to activate the immune system and aggravate liver cirrhosis^20 21^. A disordered intestinal microbiota is gradually recognized as one of the potential factors for the development of liver fibrosis. Due to the intricate ecological environment of different parts of the intestine^22^, the bacteria contained in different parts may also differ.

Ursolic acid (UA), a traditional Chinese medicine, is a natural pentacyclic triterpenoid compound derived from traditional Chinese medicine plants such as *Salvia miltiorrhiza, Hedyotis diffusa*, and *Ligustrum lucidum*, which can protect liver cells, inhibit liver inflammation and resist fibrosis^23 24^. The anti-liver fibrosis effect of UA may be related to the regulation of the activation, proliferation and apoptosis of HSCs. ^25 26^. However, the improvement effect of UA on intestinal microbiota disorder is not clear in liver fibrosis.

## METHOD

### 2.1 Experimental design and animal models

All wild-type (WT) C57BL/6 mice used in this study were from the Department of Laboratory Animal Science of Nanchang University. Mice were kept in a specific pathogen-free (SPF) environment with a 12□h light/dark cycle, a room temperature of 22±2°C, and 55±5% humidity. Mice weighing 25-35 g were randomly divided into a control group, a CCl_4_ group and a UA group (n = 8). Mice in the control group were gavaged with olive oil (2 ml/kg) twice a week for 8 weeks. Mice in the CCl_4_ group were gavaged with carbon tetrachloride (CCl_4_) (1:4 diluted in olive oil, 2 ml/kg) twice a week for 8 weeks. Mice in the UA group were gavaged with CCl_4_ for 4 weeks and then with UA (40 mg/kg/day) and CCl_4_ at the same time for the last 4 weeks. This experimental protocol meets the National Institutes of Health Guide for the Care and Use of Laboratory Animals and has been approved by the Animal Care and Use Committee of the First Affiliated Hospital of Nanchang University.

### 2.2 Histological analysis

Liver tissue was embedded in paraffin and sectioned for haematoxylin and eosin (H&E) and Masson’s trichrome staining to observe liver tissue damage and fibrosis. We randomly selected 5 visual fields of view to observe and score for liver fibrosis based on METAVIR scoring criteria.

### 2.3 Serum index test

Serum collected from mice was analysed by an automated blood biochemistry analyser to detect ALT, AST, and TBIL (Department of Clinical Laboratory, First Affiliated Hospital of Nanchang University, China).

### 2.4 Genomic DNA extraction and sequencing

Genomic DNA was extracted from the ilea and anal faeces of mice using commercial kits (Omega, China). Primers 338F 5’-ACTCCTACGGGAGGCAGCAG-3’ and 806R 5’-GGACTACHVGGGTWTCTAAT-3’ were used for polymerase chain reaction (PCR) amplification of the V3-V4 region of the 16S rRNA gene of the extracted DNA. The PCR product was sequenced using a MiSeq instrument according to the manufacturer’s guidelines (Illumina, USA).

### 2.5 Analysis of the 16S function prediction of microbiota

The 16S function prediction removes the influence of the number of copies of the 16S marker gene in the genome of the species by using PICRUSt software, which stores the COG information and KO information corresponding to the greengene id, that is, to normalize the OTU abundance table, and then to correspond to each OTU. The greengene id obtains the COG family information and KEGG Orthologue (KO) information corresponding to the OTU and calculates the abundance and KO abundance of each COG. According to the information of the COG database, the descriptive information of each COG and its functional information can be parsed from the eggNOG database to obtain a functional abundance spectrum; according to the information in the KEGG database, KO and Pathway information can be obtained, and according to the OTU abundance, the abundance of each functional category can be calculated.

### 2.6 Statistical analysis

GraphPad Prism 7.0 software was used for image production and output. Each experiment was repeated 3 times to ensure confidence in the results. A one-way analysis of variance (one-way ANOVA), Student’s t test, the Mann-Whitney rank sum test, or the Kruskal-Wallis H test was used to analyse the significant differences between groups using SPSS 23.0 software. P <0.05 was considered significant.

## RESULT1 UA reversed liver damage and fibrosis in CCl_4_-treated mice

We validated the CCl_4_-induced liver fibrosis model and the effect of UA on it. H&E and Masson’s trichrome staining on liver sections were used to determine the effect of UA on liver damage and fibrosis in CCl_4_-treated mice (Fig 1A-B). In CCl_4_-treated mice, the normal structure of the liver and the hepatic lobule was destroyed, and collagen deposition occurred, accompanied by inflammatory cell infiltration and hepatocyte necrosis. However, after UA treatment, the structure of the liver nodules and collagen deposition showed a certain improvement, accompanied by less inflammatory cell infiltration and hepatocyte swelling. Histological quantitative analysis also confirmed the improvement of liver injury and fibrosis by UA (Fig 1C-D). Moreover, we also measured serum from mice to assess liver function (Fig 1E). Compared with the control group, the serum ALT, AST, and TBIL levels in mice in the CCl_4_ group increased. The ALT, AST, and TBIL levels in mice in the UA group showed a significant decrease compared with the CCl_4_ group, indicating the protective effect of UA in liver fibrosis mice. These results suggest that UA can improve CCl_4_-induced liver damage and liver fibrosis in mice.

**Figure 1.**
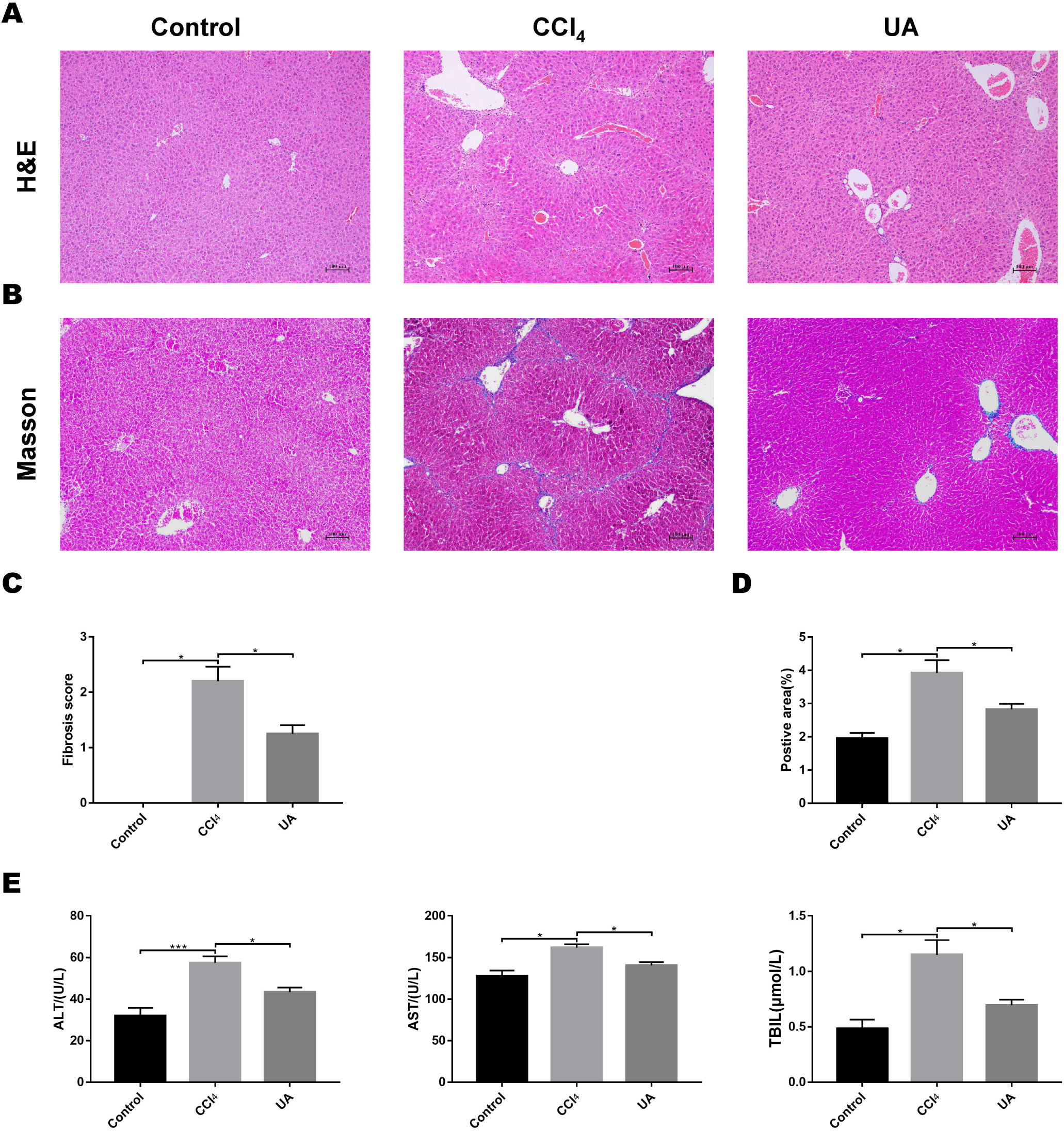
Effect of UA on liver injury and fibrosis in mice with liver fibrosis. (A) Haematoxylin-eosin (H&E) staining (100×). (B) Masson’s trichrome staining (100×). (C-D) Histological analysis of the fibrotic score and area. (E) Liver function serum index. Data represent the mean ± SD of values per group. *P < 0.05 and ***P < 0.001.

## RESULT2 High-throughput sequencing across the different anatomic sites of the mouse intestinal tract

Previous studies on the changes in microbiota in liver fibrosis often used conventional bacterial culture methods^27 28^. However, this culture technique has certain limitations, and more than 80% of the bacteria cannot be cultured^29^. As the importance of the microbiota in liver fibrotic diseases increases and more microbiota are discovered, more advanced and accurate microbial identification analysis techniques are needed. Here, we used high-throughput 16S rRNA gene sequencing technology to detect the bacteria in the ilea and faeces of the mouse model. In this study, we used a total of 2,153,777 high-quality sequences with a read length ≥200 for analysis. These high-quality sequences were assigned to 1 domain, 1 kingdom, 17 phyla, 35 classes, 66 orders, 106 families, 226 genera, and 382 species from the ileum mucosa and anal faeces of all the mice (Table 1). The rarefaction curve gradually rises until the plateau, suggesting that the sample has been fully sequenced (Fig 2A). Moreover, we also counted the number of OTUs common and unique in the sample to visually represent the similarities and differences of the microbiota composition of the samples under different treatments (Fig 2B). The results showed that in the ileal samples, there were 503 common OTUs in the control group, CCl_4_ group, and UA group and 550 common OTUs in stool samples. The total number of OTUs in the faeces is greater than in the ileum, which may suggest a higher degree of microbiota in the faeces.

**Table 1.** Bacterial communities in the ilea and faeces of mice

| Sites | Samples | Valid reads | Domains | Kingdoms | Phyla | Classes | Orders | Families | Genera | Species |
| --- | --- | --- | --- | --- | --- | --- | --- | --- | --- | --- |
| Ileum | Control | 41008 | 1 | 1 | 11 | 19 | 25 | 47 | 118 | 216 |
|  | CCl<sub>4</sub> | 35066 | 1 | 1 | 10 | 19 | 26 | 47 | 121 | 215 |
|  | UA | 34623 | 1 | 1 | 10 | 18 | 26 | 48 | 116 | 219 |
| Faeces | Control | 48914 | 1 | 1 | 14 | 26 | 55 | 92 | 169 | 269 |
|  | CCl<sub>4</sub> | 45705 | 1 | 1 | 10 | 19 | 27 | 51 | 124 | 230 |
|  | UA | 34146 | 1 | 1 | 11 | 20 | 32 | 53 | 133 | 236 |

**Figure 2.**
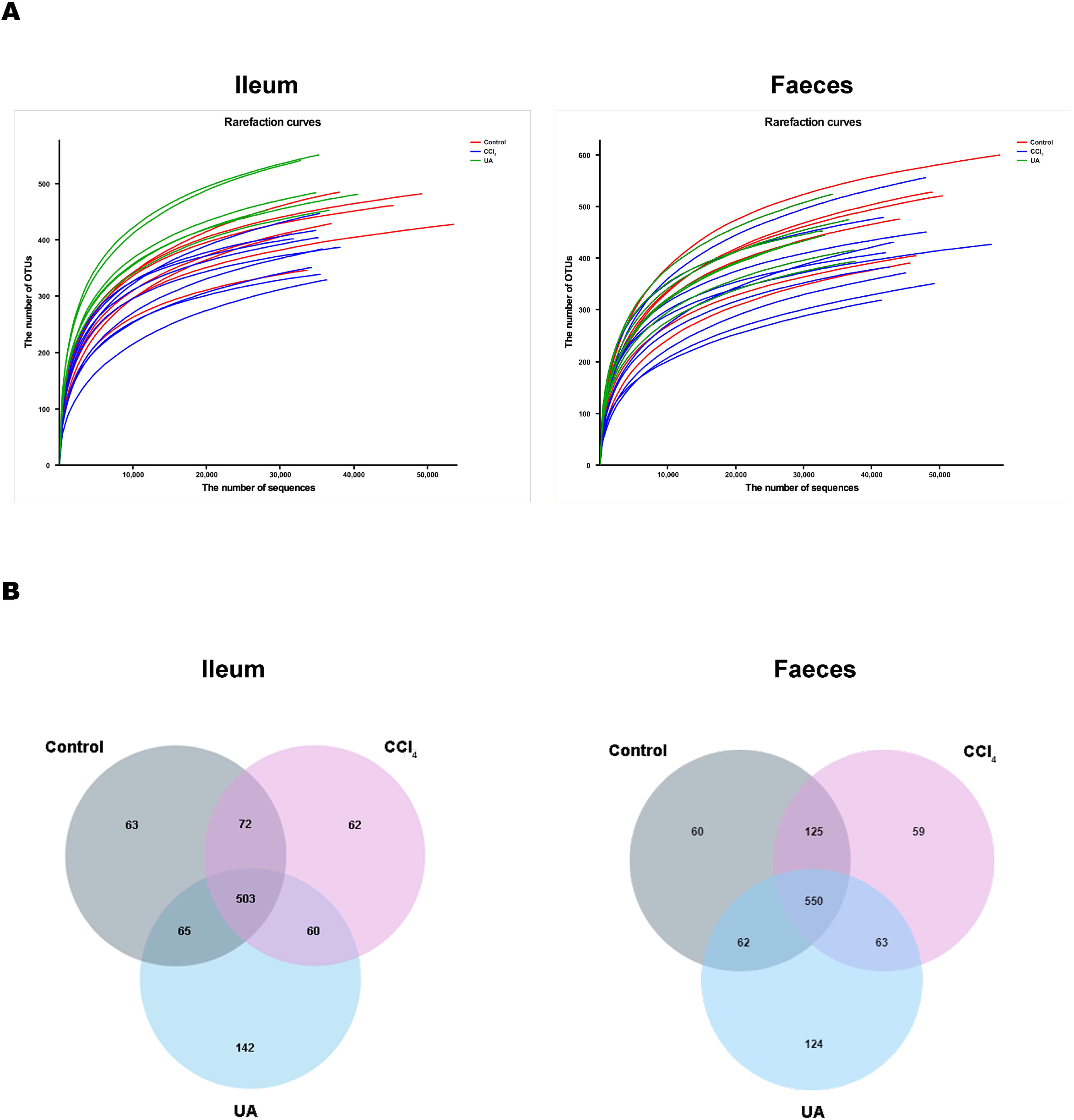
Preparation and classification of bacterial sequencing. (A) Rarefaction curves of bacterial sequencing. (B) Venn diagram showing the common and unique OTUs between different groups.

## RESULT3 E□ect of UA on the diversity of bacteria in liver fibrosis mice

To better explore the effect of UA on the bacterial microbiota of liver fibrosis, we analysed the diversity and composition of the ileal and faecal intestinal microbiota. We first tested the correlation index for the alpha diversity of the intestinal microbiota. The value of the Chao1 index, the estimated microbial abundance, showed a decrease in CCl_4_-treated liver fibrosis mice, whereas the value of this decline increased in liver fibrosis mice after UA treatment (Fig 3A). The Shannon index, used to assess the diversity of the microbiota, also showed similar changes (Fig 3B). The value of the Shannon index of bacteria in the CCl_4_ group mice was lower than that in the control group. In the UA group, the value of the Shannon index increased. In addition, the Chao1 value in the ileum was lower than that in faeces, and the Shannon index was higher in the faeces, indicating that the diversity and abundance of microbiota damage caused by liver fibrosis was partially improved after UA treatment and that the diversity of the microbiota in the ileum may be slightly higher than that in the faeces, while the abundance is the opposite.

**Figure 3.**
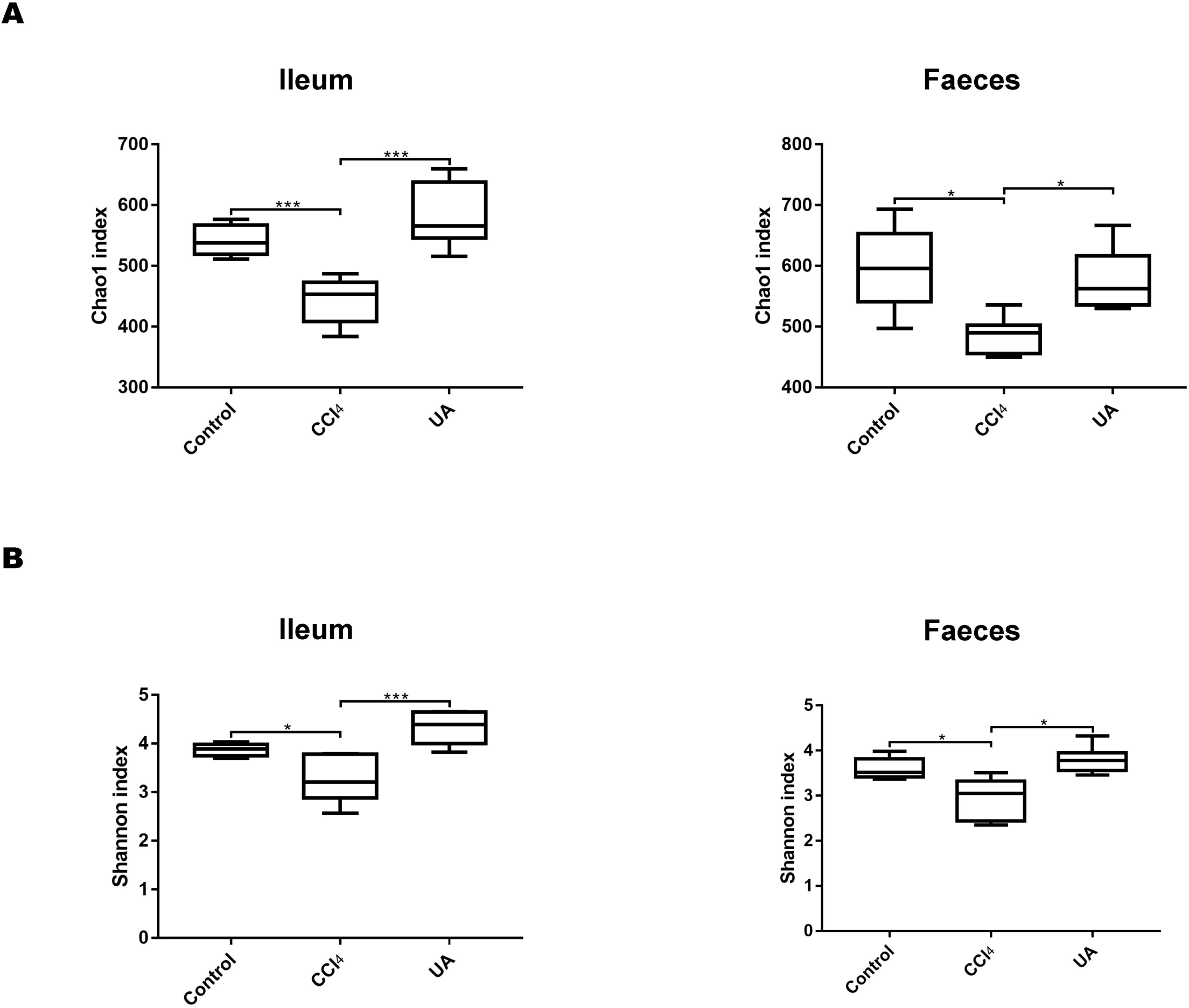
Diversity and abundance analysis of bacteria. (A) Chao1 index for assessing bacterial abundance. (B) Shannon index for assessing bacterial diversity. Data represent the mean ± SD of values per group. *P < 0.05 and ***P < 0.001.

## RESULT4 E□ect of UA on the bacterial composition of liver fibrosis mice

Liver fibrosis often affects the composition of the intestinal microbiota through the intestinal axis, further aggravating primary liver disease^27 30^. Therefore, we next analysed the changes in intestinal microbiota composition during liver fibrosis and the effect of UA on these changes. Principal coordinate analysis (PCoA) plots show that the bacterial composition of the control group, CCl_4_ group and UA group belongs to different communities (Fig 4A). We also compared the composition of the microbiota between different groups. At the phylum level (Fig 4B), Firmicutes, which includes probiotic bacteria such as *Lactobacillus*, and Bacteroidetes decrease in CCl_4_-treated mice. However, decreased Firmicutes and Bacteroidetes in CCl_4_-induced liver fibrosis mice increased after UA treatment. Elevated Verrucomicrobia in the CCl_4_ group showed a certain decrease in the UA group. At the genus level (Fig 4C), compared with the control group, Lachnospiraceae decreased in the CCl_4_ group, while *Akkermansia* increased. This change was reversed in the UA group. Moreover, linear discriminant analysis effect size (LefSe) is a software for discovering high-dimensional biomarkers and revealing genomic features. We used LefSe to reveal the composition of the microbiota of the ilea and faeces of mice in different groups (Fig 5). The results show Firmicutes enrichment in liver fibrosis mice undergoing UA treatment. However, there are separately enriched species in the ileal or faecal microbiota, indicating that UA can improve the disorder of intestinal microbiota caused by liver fibrosis, but the changes in the sites of different intestines are not completely consistent.

**Figure 4.**
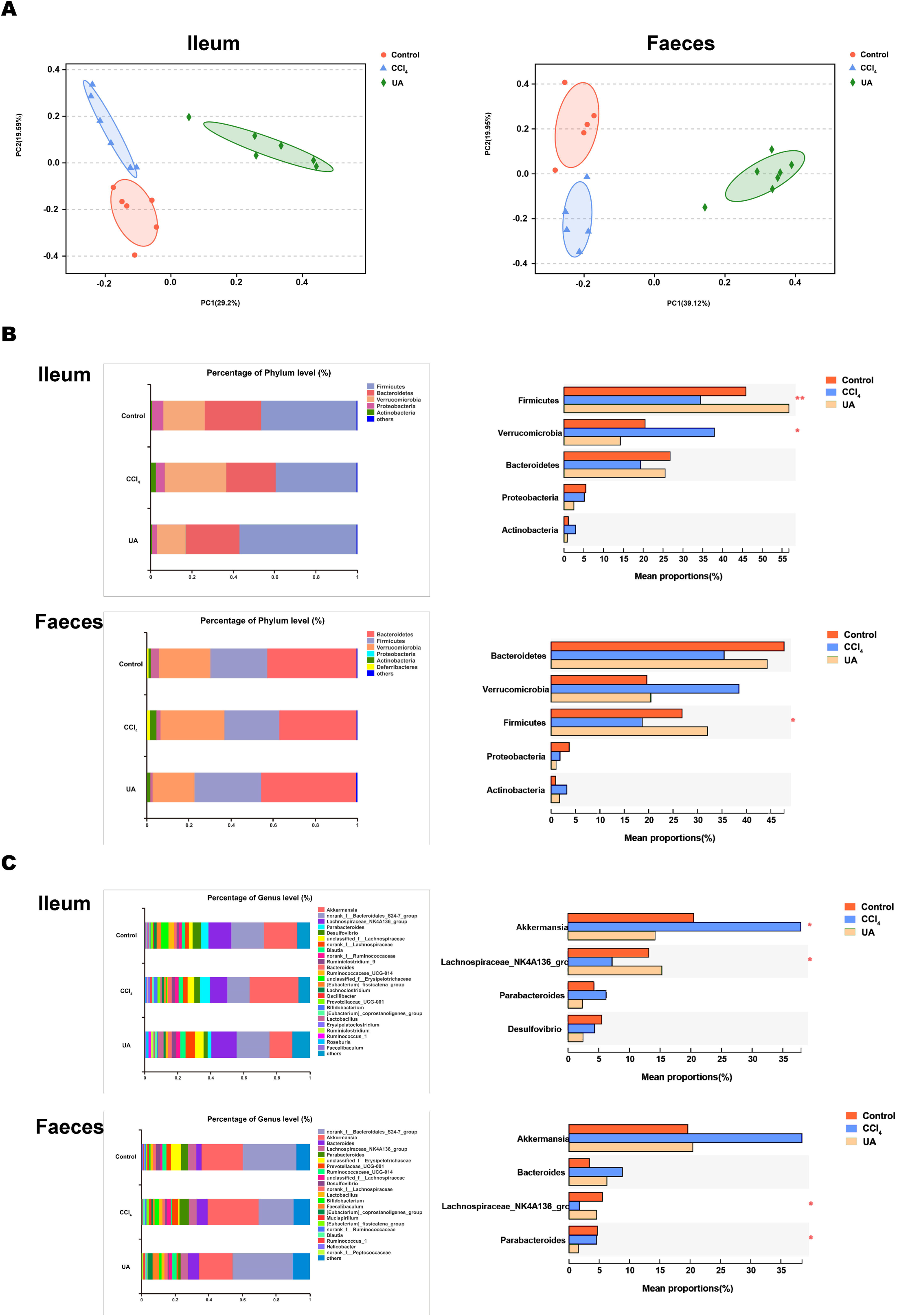
Composition analysis of microbiota. (A) Principal coordinate analysis (PCoA) plot shows the comparison of microbial community composition. (B) Bacterial composition at the phylum level. (C) Bacterial composition at the genus level. *P < 0.05 and ***P < 0.001.

**Figure 5.**
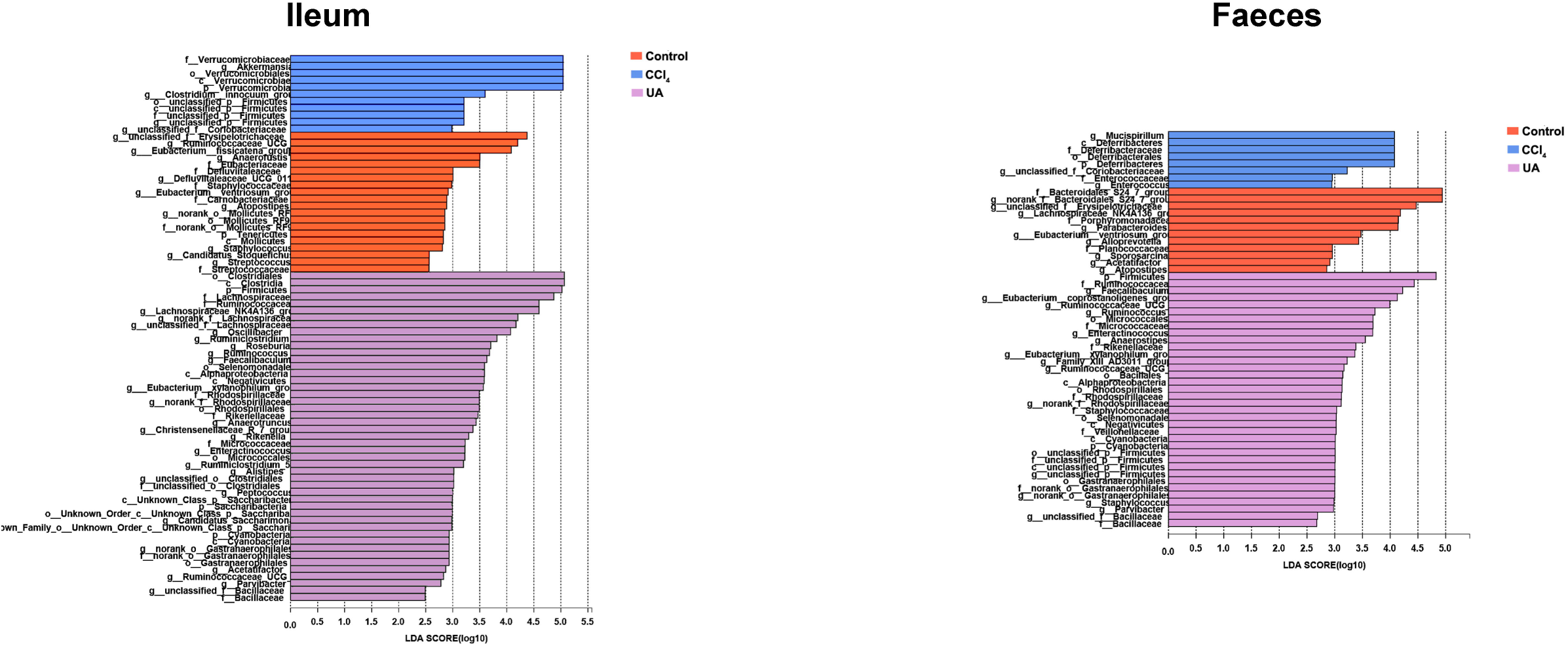
Linear discriminant analysis effect size (LefSe) for the analysis of the bacterial composition.

## RESULT5 Relationship between microbial community structure and biochemical factors

Biochemical indicators in the liver during liver fibrosis often change abnormally^31^. These indicators can partially reflect the development of liver fibrosis. However, the link between these indicators and the gut microbiota has not been confirmed. To this end, we used distance-based redundancy analysis (db-RDA) to compare microbial species with environmental variables to clarify the relationship between serum indicators and the bacterial community. The results suggest that in the ileum, ALT (P = 0.001), AST (P = 0.002), and TBIL (P = 0.001) can significantly affect the bacterial community in the ileum. The results of the microbiota in the faeces are similar. ALT (P=0.001), AST (P=0.002), and TBIL (P = 0.005) significantly affected the bacterial structure (Fig 6A). To further clarify the effects of these serum indicators on the composition of the intestinal microbiota, we completed a correlation heatmap analysis. As shown in Fig 6B, ALT was significantly positively correlated with Actinobacteria (P=0.038) and Erysipelotrichia (P = 0.01). AST was significantly positively correlated with Erysipelotrichia (P = 0.003) and Mollicutes (P = 0.019) in faeces. In the ileum, ALT and AST are positively correlated with Gammaproteobacteria, and TBIL is positively correlated with Betaproteobacteria, but it is not statistically significant.

**Figure 6.**
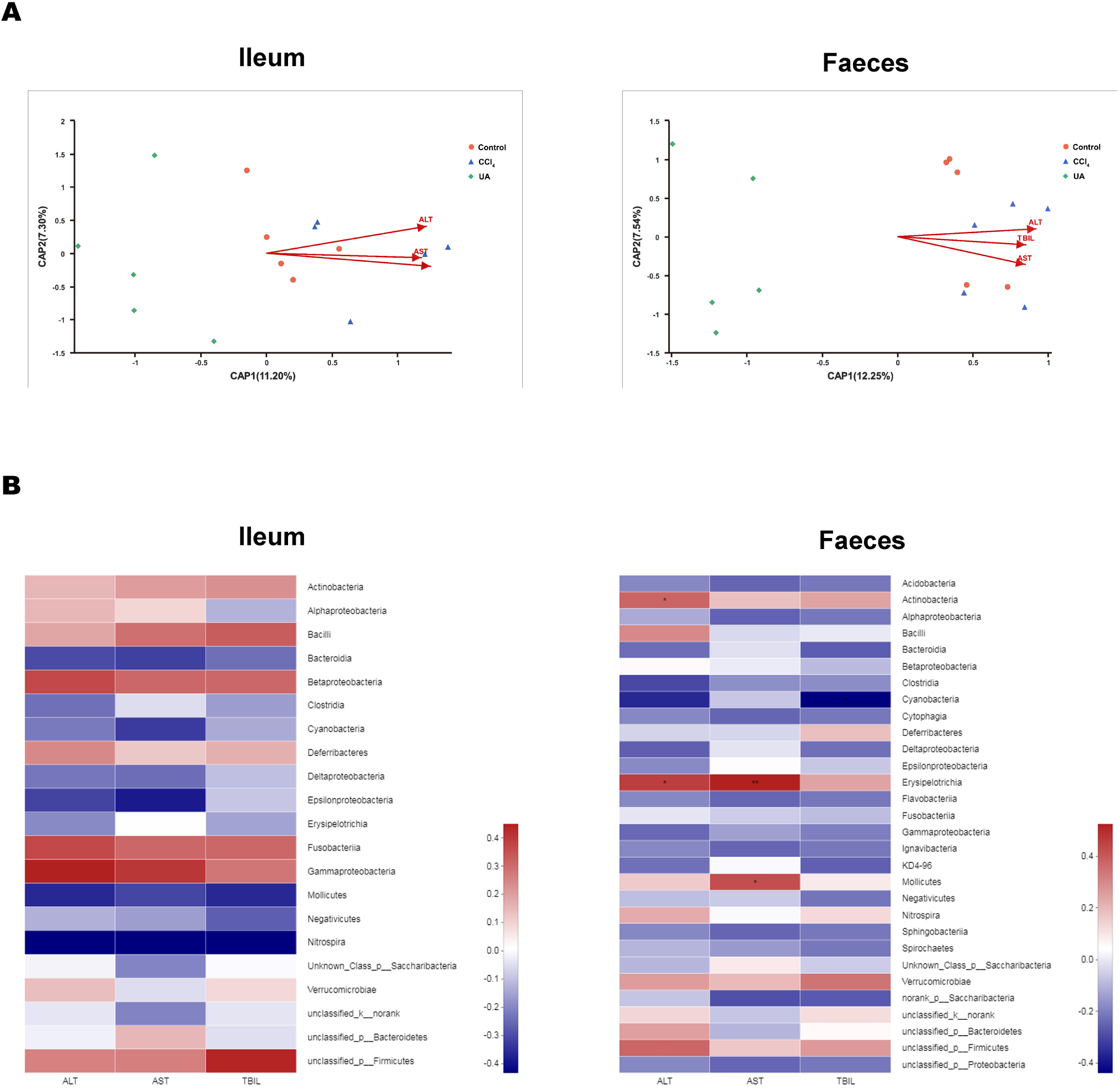
Correlation analysis between bacteria and serum indicators. (A) Distance-based redundancy (db-RDA) analysis is used to assess ALT, AST, and TBIL and bacterial structure correlations. (B) Heat mapping is used to assess the correlation between ALT, AST, and TBIL and bacterial species. *P < 0.05 and ***P < 0.001.

## RESULT6 Prediction and analysis of microbiota function

Changes in the microbiota under different body conditions indicate some changes in metabolic function. We predicted the function of microbial changes in different animal models by querying the COG and KEGG databases. The COG database included pathways for bacterial and archaeal genomes. Compared with control mice, the enriched pathways were “Cell wall/membrane/envelope biogenesis”, “Intracellular trafficking, secretion, and vesicular transport”, and “Secondary metabolites biosynthesis, transport and catabolism” in the bacteria of the ilea of mice in the CCl_4_ group. After UA treatment, enriched pathways were transformed into “Carbohydrate transport and metabolism”, “Transcription”, “Defense mechanisms”, and “Translation, ribosomal structure and biogenesis” (Fig 7A). In faeces bacteria, liver fibrosis mice enriched pathways were “Intracellular trafficking, secretion, and vesicular transport”, “Signal transduction mechanisms”, and “Secondary metabolites biosynthesis, transport and catabolism”. The enriched pathways of faeces in UA-treated liver fibrosis mice were “Translation, ribosomal structure and biogenesis”, “Transcription”, “Defense mechanisms”, “Replication, recombination and repair” and “Energy production and conversion” (Fig 7B).

**Figure 7.**
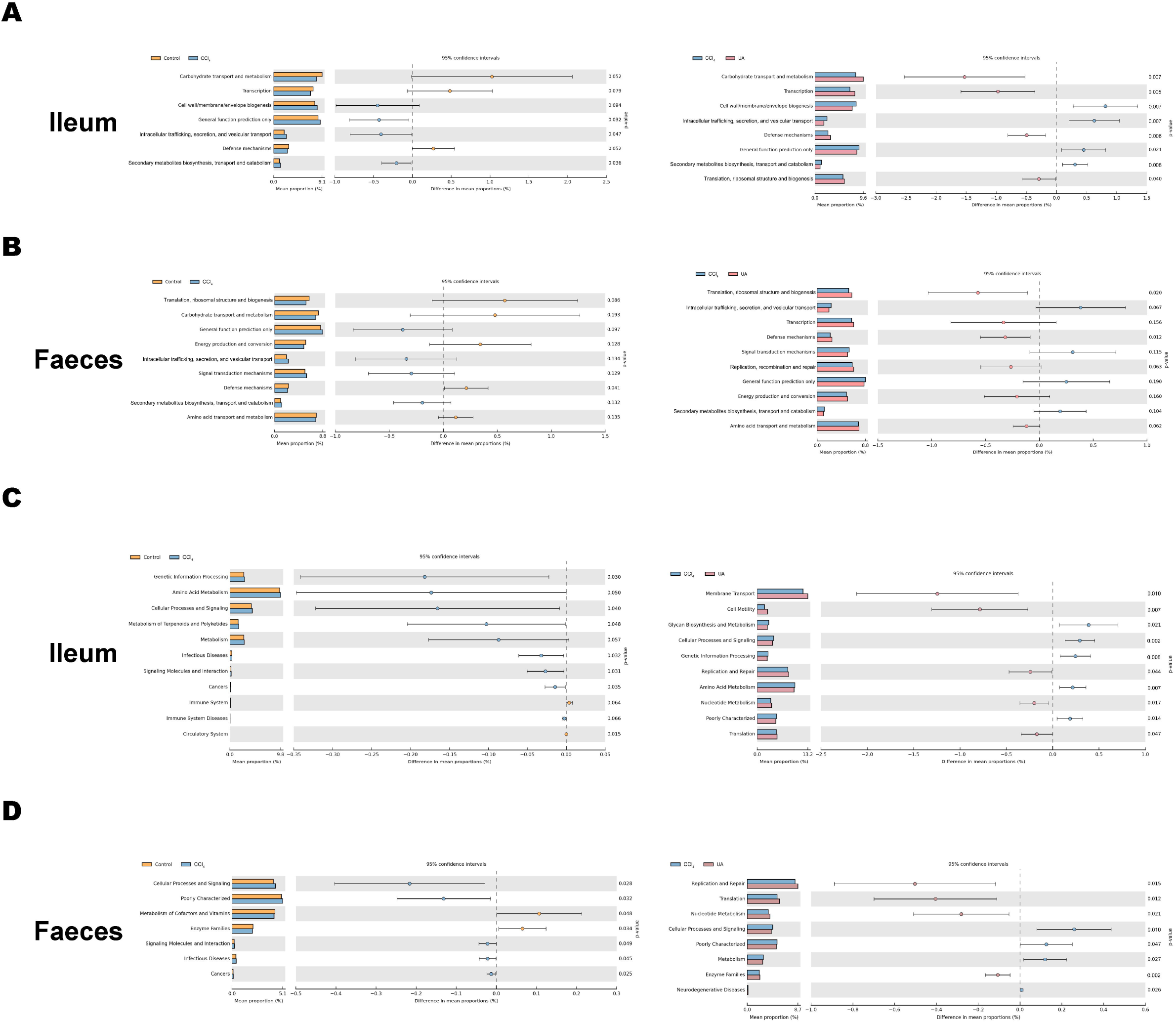
Functional prediction analysis of microbiota. (A) COG analysis of major enrichment and differential pathways in all groups in the ileal bacteria. (B) COG analysis of major enrichment and differential pathways in all groups in the faecal bacteria. (C) KEGG analysis of major enrichment and differential pathways in all groups in the ileal bacteria. (D) KEGG analysis of major enrichment and differential pathways in all groups in the faecal bacteria.

The KEGG database mainly contains information about metabolic signalling pathways. We performed functional predictions on the microbiota of different parts by searching the KEGG database. In the bacteria of the ilea of mice in the CCl_4_ group, the main metabolic KEGG terms were “Genetic Information Processing”, “Amino Acid Metabolism”, and “Cellular Processes and Signaling”. Compared with the CCl_4_ group, the main KEGG terms were “Membrane Transport”, “Cell Motility”, “Replication and Repair”, “Nucleotide Metabolism”, and “Translation” in the bacteria of the ilea of mice in the UA group (Fig 7C). Then, in the functional prediction of faecal microbiota, CCl_4_-induced liver fibrosis mice mainly included the metabolic pathways “Cellular Processes and Signaling”, “Poorly Characterized”, and “Signaling Molecules and Interaction”. The main metabolic pathways in UA-treated liver fibrosis mice were “Replication and Repair”, “Translation”, “Nucleotide Metabolism”, and “Enzyme Families”, similar to the control mice (Fig 7D).

## DISCUSSION

In this study, we show that UA can improve the disorder of intestinal microbiota during liver fibrosis. Based on the latest generation of high-throughput sequencing technology, we have investigated the changes in the intestinal microbiota across different anatomic sites of the mouse intestinal tract.

Due to the presence of the liver-gut axis, the disorder of normal metabolic activity in the liver during chronic liver disease can cause damage to the intestinal homeostasis ^32 33^. Therefore, changes in the intestinal microbiota have been found in an increasing number of chronic liver diseases^34-36^. However, there are few studies on changes in intestinal microbiota during liver fibrosis. Here, we analysed the changes in intestinal microbiota during liver fibrosis. In CCl_4_-induced liver fibrosis mice, the diversity and abundance of the microbiota showed a significant decrease compared with normal mice. An analysis of the composition of the microbiota showed that the beneficial bacteria in the liver fibrosis mice also decreased. Functional predictions suggest that this change in the microbiota in liver fibrosis may indicate the occurrence of adverse pathways, suggesting that there is a disorder in the intestinal microbiota during liver fibrosis.

The outstanding findings in this paper are that the anti-liver-fibrosis Chinese medicine UA also has a certain improvement effect on the intestinal microbiota disorder during liver fibrosis. Our group has previously confirmed that UA can protect liver tissue and reverse liver fibrosis in an animal model of liver fibrosis induced by CCl_4_^37^. In addition, our previous experiments also found that UA can reduce intestinal damage and protect intestinal integrity during liver fibrosis. Therefore, we speculate that UA improves intestinal microbiota disorders in liver fibrosis mice. In the liver fibrosis mice that underwent UA treatment, the diversity and abundance of the intestinal microbiota showed a significant increase. Similarly, the proportion of beneficial bacteria in UA-treated mice also increased. Functional predictions indicate the occurrence of favourable pathways. The microbiota of liver fibrosis mice after UA treatment approached the microbiota of normal mice. These results confirm our hypothesis that UA can improve the disorder of intestinal microbiota during liver fibrosis. This may be the potential mechanism for UA to exert its anti-fibrosis effects, that is, to improve the disordered microbiota and to achieve the reversal of liver fibrosis through the liver-gut axis.

The intestine is an important part of the digestive system and the longest part of the digestive tract^38^. The ecological environment of different regions of the intestine is inconsistent^22^, which may affect the ecology of the microbiota contained here. Previous studies on the intestinal microbiota of liver fibrosis have often only studied and analysed the microbiota of one site. In this study, the intestinal microbiota of the ileum and faeces were collected for analysis to better define the changes in intestinal microbiota during liver fibrosis and the effect of UA on this change. The results also confirmed that there are some differences between the microbiota in the ileum and the microbiota in the faeces. Interestingly, in the ileal microbiota, the Bacteroidetes is lower than in the faecal microbiota. This may be related to the presence of more oxygen in the ileum, which is not conducive to the growth of anaerobic bacteria. Based on this finding, studies related to microbiota should pay attention to the location of the intestinal collection of the microbiota, which is of great significance for the study of the relationship between the microbiota and the development of liver fibrosis.

In conclusion, our research indicates that intestinal microbiota are disordered during liver fibrosis and that UA has an effect on the disorder. This provides a new perspective for revealing the potential mechanism of UA to reverse liver fibrosis. However, the molecular mechanism and clinical data of UA to improve intestinal microbiota are still lacking, and subsequent improvement is needed. This study provides theoretical support for the future use of UA in the clinical targeting of the intestinal microbiota for the treatment of liver fibrosis.

## Author Contributions

SW designed and wrote the manuscript. CH analysed the data. SW critically revised the manuscript.

### Funding

This study was supported by the National Natural Science Foundation of China (grant number: 81960120) and the “Gan-Po Talent 555” Project of Jiangxi Province.

## Acknowledgements

We would like to thank the National Natural Science Foundation of China for economic support.

